# Heterogeneous run-and-tumble motion accounts for transient non-Gaussian super-diffusion in haematopoietic multi-potent progenitor cells

**DOI:** 10.1101/2021.11.19.469302

**Authors:** Benjamin Partridge, Sara Gonzalez Anton, Reema Khorshed, George Adams, Constandina Pospori, Cristina Lo Celso, Chiu Fan Lee

## Abstract

Multi-potent progenitor (MPP) cells act as a key intermediary step between haematopoietic stem cells and the entirety of the mature blood cell system. Their eventual fate determination is thought to be achieved through migration in and out of spatially distinct niches. Here we first analyze statistically MPP cell trajectory data obtained from a series of long time-course 3D *in-vivo* imaging experiments on irradiated mouse calvaria, and report that MPPs display transient super-diffusion with apparent non-Gaussian displacement distributions. Second, we explain these experimental findings using a run-and-tumble model of cell motion which incorporates the observed dynamical heterogeneity of the MPPs. Third, we use our model to extrapolate the dynamics to time-periods currently inaccessible experimentally, which enables us to quantitatively estimate the time and length scales at which super-diffusion transitions to Fickian diffusion. Our work sheds light on the potential importance of motility in early haematopoietic progenitor function.

---

The haematopoietic system is responsible for the generation of billions of new blood cells daily. This considerable feat is orchestrated from the bone marrow whose constituent blood cells are organized into a hierarchical lineage tree, atop of which reside haematopoietic stem cells (HSCs). HSCs have two defining properties; self-renewal - the ability to replenish their own numbers, and multi-potency - the potential to differentiate into any given blood cell type. Downstream of HSCs lie multi-potent progenitor cells (MPPs), which act as an intermediary between HSCs and mature blood cells. Their successive proliferation and differentiation amplifies cell numbers to enable a small pool of HSCs to achieve a staggering output of mature blood cells [1].

To achieve tissue homeostasis and to properly respond to stress and infection cell division and differentiation must be tightly regulated. The motility of haematpoietic stem and progenitor cells (HSPCs) is emerging as a crucial component to these regulations [2]. Therefore, a detailed understanding of the spatial organization and dynamics underpinning the relationship between HSPCs and their environment is crucial. Intravital microscopy has been a key experimental technique that directly probes this relationship [3, 4]. This is of special clinical interest in the irradiated state, due to the importance of radiation therapy in conditioning the bone marrow prior to transplantation when treating malignancies of the blood.

In parallel development, the diffusive properties of migrating organisms have been the subject of intense interest within the statistical physics community [5, 6]. A common characteristic amongst such systems is a non-linear, power-law scaling of the mean square displacement (MSD) with time. This phenomenon stands in contrast to the linear relationship expected in the ‘typical’ case of Fickian diffusion. This ‘anomalous’ diffusion - described as super-diffusion (sub-diffusion) in cases where the MSD growth exceeds (falls below) the linear case - has been postulated to influence a variety of biological processes spanning virtually all biological length-scales [7–21]. Two recent intravital imaging studies have examined the dynamical behavior of HSPCs in their native, steady-state environment and found that i) progenitor cells display enhanced motility relative to HSCs, and ii) temporal heterogeneity within HSC trajectories, where the dynamical behavior alternated between periods of a confined random walk and processive motion [2, 4]. There are a limited number of prior works investigating the migratory behavior of transplanted HSCs [22, 23], and to our knowledge a quantitative analysis of the diffusive properties of haematopoietic MPPs in an irradiated setting has yet to have been undertaken.

In this article, we first report on a statistical analysis of cell trajectory data taken from 3D *in-vivo* imaging experiments of haematopoietic multi-potent progenitor cells in the irradiated bone marrow cavity of murine calvaria. Many of the cell trajectories are observed over long time periods atypical of 3D *in-vivo* bone marrow imaging experiments, with 44% of trajectories having a length greater than 3 hours and 17% having a length of greater than 6 hours. We demonstrate that the cells display transient non-Gaussian super-diffusion over time-scales of biological interest. We then explain this observation using a data-driven run-and-tumble (RTM) model which takes into account heterogeneity in the dynamics of the ensemble. We found that the incorporation of heterogeneity into our RTM is necessary and sufficient to explain the non-Gaussian super-diffusive behavior. We extrapolate the dynamics to time-periods currently inaccessible to time-lapse imaging experiments which enables us to quantitatively estimate the time and length scales at which super-diffusion transitions to Fickian diffusion. These estimates will be integral to understanding how stem and MPP cell motility influence the regulations of blood cell generation due to the recognized importance of spatial organization in controlling the function of the haematopoietic stem cell niche.

## MULTI-POTENT PROGENITOR CELLS DISPLAY NON-GAUSSIAN SUPER-DIFFUSION

### Experimental Data

The cell trajectory data presented were extracted from a series of *in-vivo* imaging experiments in which labeled multi-potent progenitor cells expressing membrane-bound green fluorescence protein (GFP) purified from the bone marrow of donor mice are transplanted into myeloablative recipient mice and their three-dimensional positions followed using confocal microscopy two days after radiation therapy for time periods ranging from 18 to 525 minutes with three-minute intervals between subsequent frames (see figure S1 in [37]). In figure 1(a) we show a maximum *z*–projection still frame depicting a typical configuration of such an experiment with a few haematopoietic MPP cells (green) and the local micro-environment. A representation of an extracted MPP trajectory is shown in figure 1(b).

**FIG.1.**
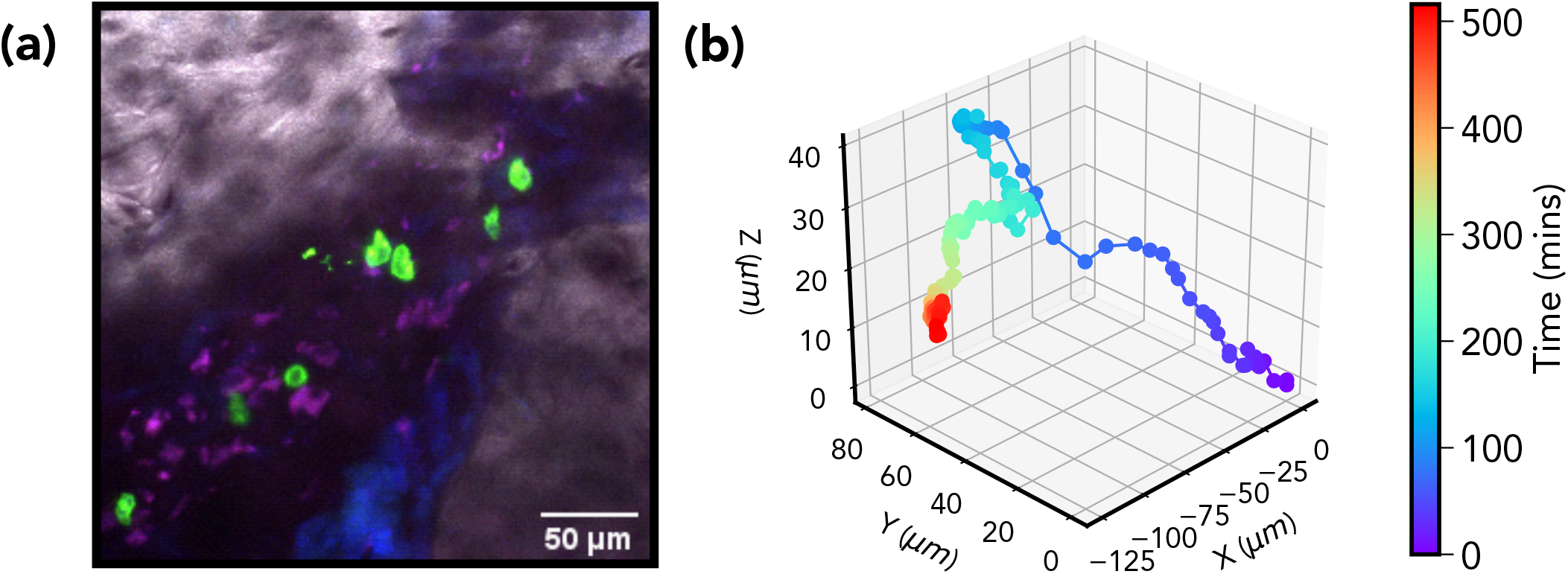
*Experimental Data* - (a) Haematopoietic multi-potent progenitor cells (green) and the local micro-environment in an irradiated bone marrow cavity taken from a time-lapse in-vivo imaging experiment (grey: bone; blue and purple: autofluores- cence). (b) 3D reconstruction of a MPP trajectory extracted from the same set of experiments.

### Anomalous Diffusion

The standard indicator of anomalous diffusion is a power-law scaling of the mean square displacement

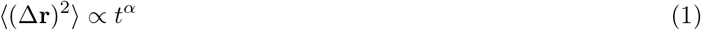

where the angular brackets indicate an ensemble average. An exponent of *α* > 1 indicates super-diffusion, which is the regime of interest here. The underlying mechanism responsible for this result depends upon the nature of the stochastic process generating the motion. For instance, a continuous-time random walk in which the displacement of random walker moving at a fixed speed is given by

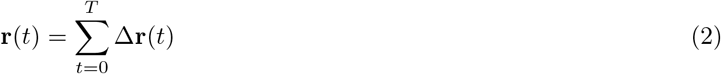

where Δ**r** are run lengths represented by random variables distributed according to *P* (Δ**r**, *t*). In the limit of large *T*, assuming independent run lengths with finite variance, the Central Limit Theorem (CLT) dictates that the distribution Prob(**r**, *t*) of the ensemble (also known as the propagator) will converge to a Gaussian distribution with a second moment that scales linearly with time. Super-diffusion occurs when the assumptions of the CLT are violated. Models with heavy-tailed step length distributions, such as Levy walks [24], violate the finite variance assumption. Whilst models with persistent correlation, such as the Elephant walk [25, 26], violate the assumption of statistically independent displacements. Furthermore, a number of studies [9, 11, 12] have shown empirically that a useful investigative tool is the scaling property

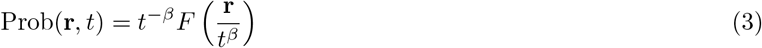

where *F* is generally non-Gaussian and *β >* 0.5.

Due to the limited amount and length of trajectory data typically collected during *in-vivo* imaging experiments direct access to the ensemble averaged MSD is not possible. Thus in order to improve the statistical properties of the estimator the time-averaged mean square displacement is computed instead (TAMSD) [5].

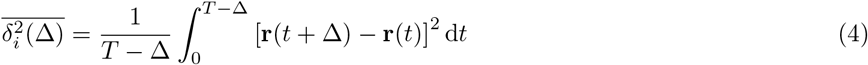

The subscript *i* indicates the individual cell in question and the over-script bar the time average. This quantity may then be further averaged over all *N* tracks

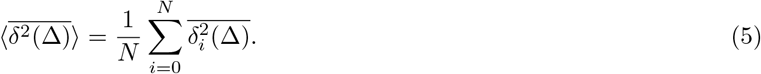

In figure 2 we present the evidence for transient MPP super-diffusion, 2(a) shows the the power-law scaling (plotted in log-log scale) of the TAMSD averaged over all cells. In this figure we have taken a maximum lag-time of two and a half hours, and to ensure a reasonable statistical convergence for later time points we have used tracks of three hours or more in length. An ordinary least-squares fit yields a gradient of *α* = 1.22, indicative of a super-diffusive scaling.

**FIG.2.**
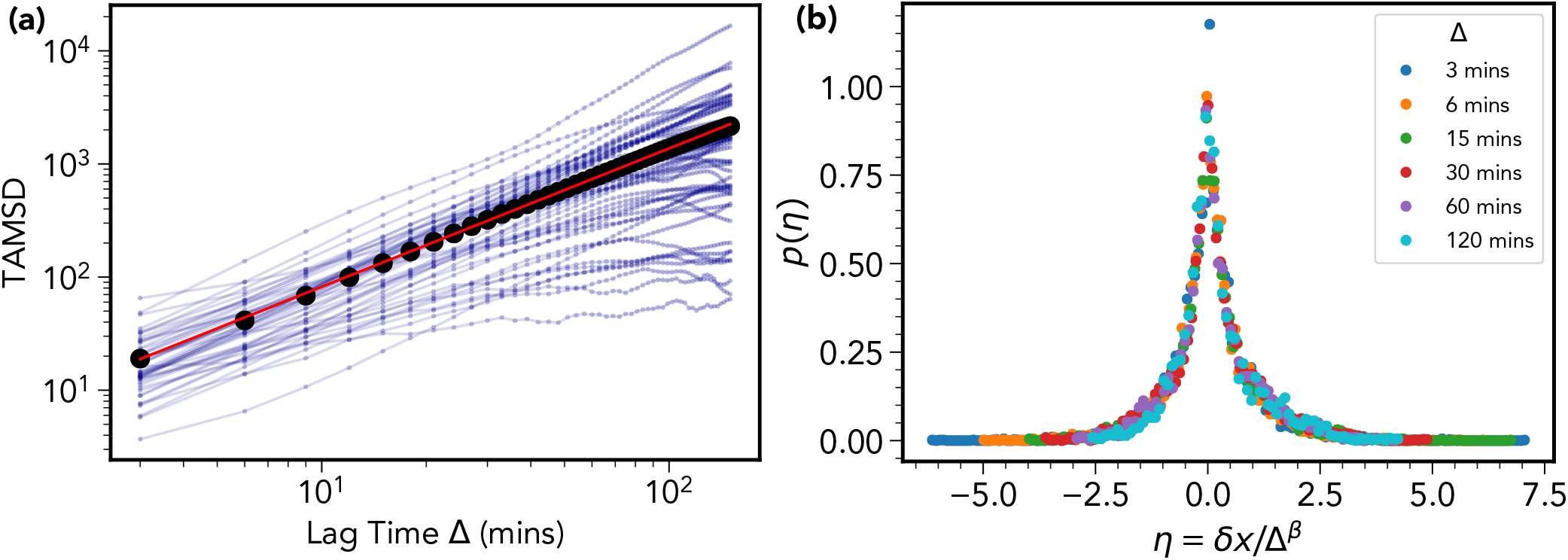
*Multipotent progenitor cells display transient non-Gaussian super-diffusion* - (a) Time-averaged mean square displacement trajectories for each cell of track length greater than 3 hours (blue curves) along with the average over all cells (black curve). A linear fit (red curve) yields a super-diffusive exponent of *α* = 1.22. (b) Curve collapse of the distribution of the re-scaled variable *η* = *δx/*Δ^*β*^, where *δx* = *x*(*t* + Δ) − *x*(*t*), for 6 different lag time intervals Δ. The value of the exponent *β* is estimated to be 0.72 using a procedure described in the SI. For standard Fickian diffusion it is expected that *α* = 1 and *β* = 0.5.

In 2(b) we plot the probability distributions of the re-scaled variable *η* = *δx/*Δ^*β*^ for various lag times Δ, where *δx* = *x*(*t* + Δ) − *x*(*t*); − *x*(*t*) being the *x* -coordinate of a given cell at time *t*. The value of the exponent *β* was determined to be 0.72 using a procedure described in the SI (see figure S2 in [37]). A similar result was observed for the *y*-coordinate, however, the *z*-coordinate showed a lesser value of 0.59 - this is likely an artifact of the limited field of view in the *z*-direction, a common limitation of 3D intravital imaging experiments. The resulting curve-collapse demonstrates the super-diffusive scaling property expressed in equation 3 over a three hour time window. It is also of note that shape of the distributions appears to be distinctly non-Gaussian, a characteristic also quantified in [37] (see figures S3 and S4).

## RUN AND TUMBLE MODEL EXPLAINS TRANSIENT SUPER-DIFFUSION

To explain the apparent transient super-diffusion displayed by the MPP population we implement a run and tumble (RTB) model of cell motility. RTB models have been successful in the dynamical description of a variety of motile organisms including bacteria *E. coli* and Salmonella [27, 28], and the eukaryotic unicellular alga Chlamydomonas [29, 30]. A random walker undergoes intermittent linear ‘runs’ of length *r* for a time-interval Δ*t* before re-orientating its direction by angle *θ* and proceeding to embark on another run. In the simplest case *r* and *τ* are related through a fixed speed *V*. This case would not provide a faithful representation of our data-set due to internal heterogeneity within the cell trajectories. Our model requires the specification of the run length distribution *f* (*r*), the turn angle distribution *h*(*θ*), and the run time distribution *g*(Δ*t*).

To avoid the arbitrary imposition of a parametric model onto the data-set we take a data driven approach, in which the relevant distributions specifying the model correspond to empirical distributions constructed from the MPP data-set. This allows for the avoidance of model error in our inference. We discuss the details of our approach in the SI, while we briefly describe our procedure here. The empirical distributions are constructed by coarse-graining each trajectory to include points with a threshold value of at least 7.2 microns (approximately one cell diameter) separation. The threshold value was determined as described in the SI. This procedure is done to remove short-range, transient fluctuations in the cell centroid position unrelated to translocations of the entire cell body. Small changes to the threshold value do not significantly alter the results [37] Displacements between points on these coarse-grained trajectories enable the definition of run length, run time, and turn angle distributions. In the SI, we further implement the more elaborate Bayesian method of [31] to complement and to confirm the results of the minimal RTB model presented herein.

We simulate a trajectory using the following procedure which is described in detail in the supplementary information:

1. Assign average run length 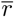, run time 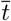 and average turn angle 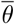.
2. Draw from scaled run length distribution 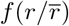 and multiply by 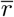 to obtain actual run-time.
3. Draw from scaled turn angle distribution 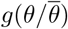 and multiply by 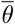 to obtain actual turn angle.
4. Draw from scaled run time distribution 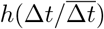 and multiply by 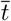 to obtain actual run-time.
5. Update time and position variables and return to 2.

Note that, our model incorporates the observed inter-track heterogeneity. This is manifested via significant inter-trajectory variation in the mean run length and run-time and mean turn angle. We account for this by drawing from scaled empirical distributions. For example, each experimental run-time interval is scaled by the average for the corresponding cell trajectory. These dimensionless values are then aggregated across all trajectories to obtain the model empirical distribution functi<u>on</u>. In other words, for each trajectory we h<u>ave</u> a set of run times 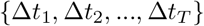 and an associated mean run-time 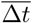. For each simulated track we assign a 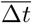 from the list of empirical mean runtimes and then generate (dimensionless) samples from the distribution 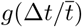. We then obtain a true run-time in minutes through multiplication by the corresponding scale factor 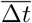. Run length and angular heterogeneity was incorporated by applying the same procedure. Heterogeneous diffusion processes have been implicated in the emergence of non-Gaussian displacement distributions [31–35]. Figure 3 shows that our data-driven model accounts for the super-diffusive behavior observed in the data. In the SI we demonstrate that the ensemble of simulated trajectories display non-Gaussian displacement distributions similar to those computed from the data.

**FIG.3.**
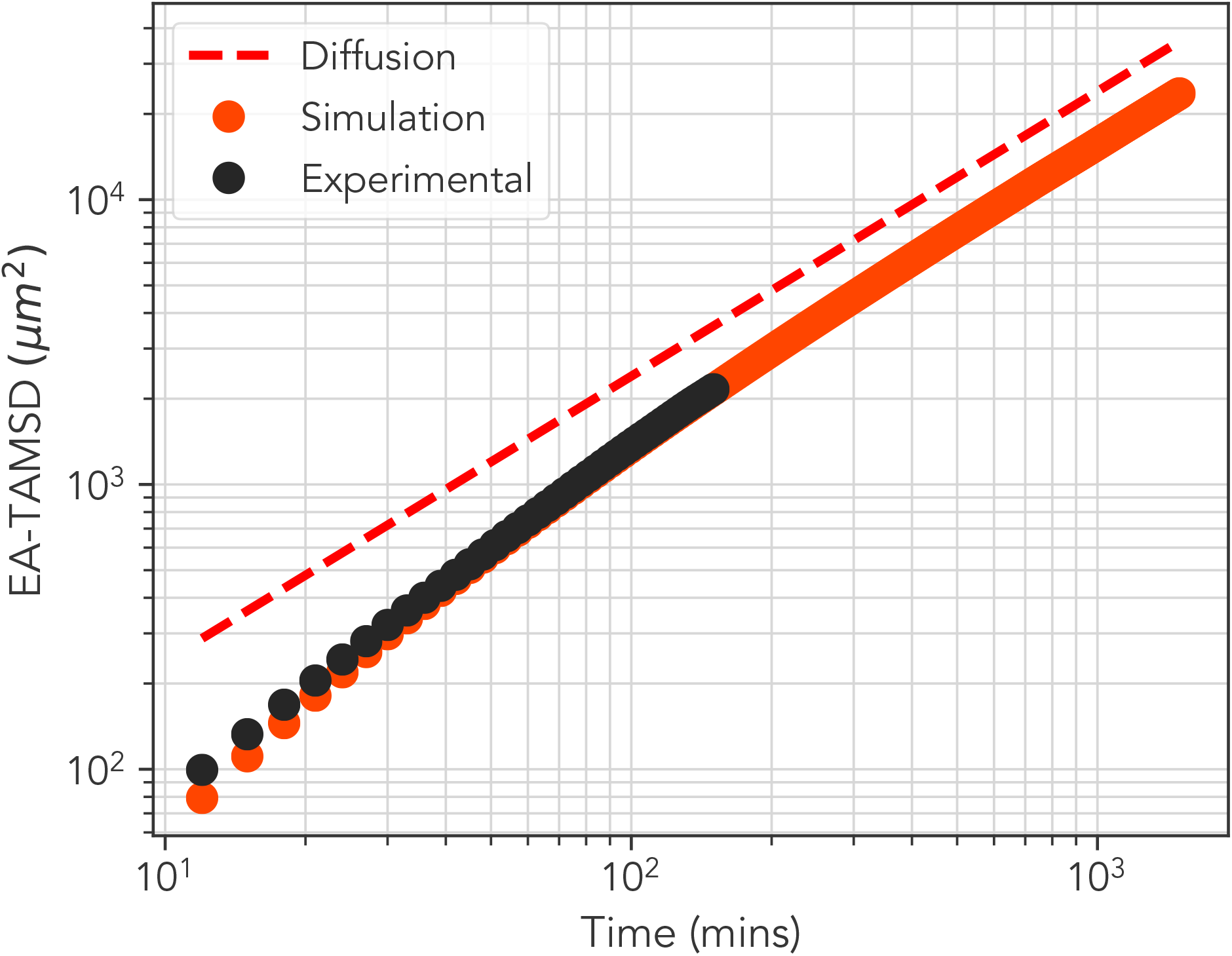
*Comparison of simulation results and experimental data* - Ensemble averaged TAMSD curves for experimental (black) and simulation (orange) alongside the line of diffusion (red dashed), which is included as a guide to the eye. The simulation data has been extended into a region inaccessible to experimentation in which a crossover to diffusion is observable. Error bars are too small to be visible.

Beyond explaining the experimental findings, our simulation model also enables us to investigate MPP dynamics at timescales inaccessible to intravital microscopy experiments. The curve is the result of simulating 51 trajectories, one for each average run-time calculated from the trajectory data, for 30000 minutes, and the average taken using a maximum lag-time of 1500 minutes, an order of magnitude greater than the empirical data. We can infer by inspection of this result that the transition to Fickian diffusion takes place at around the 100-200 minute mark. If we refer to the most exploratory cell in 2(a) we can then infer an approximate upper limit to the spatial region covered by the persistent motion to be at least 100*µm* or around 20 cell diameters in length.

## BIOLOGICAL IMPLICATIONS OF OUR RESULTS

We have demonstrated that the haematopoietic multi-potent progenitor cells display transient super-diffusion which can be explained by a run-and-tumble model incorporating one necessary and sufficient feature: heterogeneity between cell tracks.

Our results did not show any evidence to suggest that the transient super-diffusion observed in the MPP trajectory data could be attributed to heavy-tailed run length distributions, precluding a Levy Walk style model. Instead, incorporation of heterogeneity in run-length, turn-angle, and run-time sufficiently re-capitulated the experimental TAMSD.

Our model-facilitated extrapolation point to an upper limit on the range of influence of the directed motion of MPPs within the bone marrow cavity of irradiated murine calvaria. We infer this limit from the cross-over to Fickian diffusion of the simulated TAMSD. The high value of the upper limit is surprising in part because it spans several multiples of cell diameter, suggesting that persistent motion may play a significant role in determining the bio-physical properties of early stage bone marrow tissue regeneration. A key concept within haematopoietic stem cell biology is that of a niche, a distinct anatomical compartment within the bone marrow whose cellular constituents directly regulate cell fate. Whilst our study makes no direct connection between the bone marrow micro-environment and MPP motility, it is reasonable to suggest that the persistent motility observed may serve as a stochastic mechanism through which progenitor cells are able to re-locate to regions within close proximity to potential niches.

The heterogeneity present within the cell motion is notable, and in line with the phenotypic and functional heterogeneity observed in the biological literature [36] - there is significant variation in the average run-time and turn-angle for each cell, evidently giving rise to a spectrum of diffusivities. This may be attributable to differing transcriptional states in the genes associated with motility present within the population of cells.

In summary, we have employed a data-driven RTM to explain the non-Gaussian, super-diffusive dynamics of MPP observed in long time-course 3D *in-vivo* imaging experiments on irradiated murine bone marrow, and used our model to quantify the temporal and spatial ranges in which these anomalous features are present. Interesting future directions will be to connect these anomalous dynamics to bone marrow regeneration, niche organization, and homeostasis.

