## Supplementary Information for "Heterogeneous run-and-tumble motion accounts for transient non-Gaussian super-diffusion in haematopoietic multi-potent progenitor cells"

### EXPERIMENTAL DESCRIPTION

Bone marrow 3D intravital imaging experiments aim to selectively track the positions of cells in a given subset of the haematopoietic lineage tree in order to observe their dynamical behavior. Such information is crucial to a complete understanding of the processes which underpin haematopoietic stem and progenitor cell (HSPC) biology; allowing insight into cell motility, cell division and apoptosis, and their relation to the bone marrow micro-environment.

Haematopoietic stem cells (HSCs) and multipotent progenitor cells (MPPs) are defined operationally through their ability to provide long- and sort-term reconstitution capacities, respectively, to the haemopoietic tissue. However, identifying HSCs using such a definition requires the use of serial transplantation assays, clearly limiting the ability to perform experiments to visualize HSCs and related populations *in-vivo*. Instead, haematopoietic cell populations are identified phenotypically using cell surface marker proteins. For example, haematopoietic progenitors may be identified by selecting populations of cells lacking the expression of a cocktail of cell surface markers associated with terminal differentiation - termed lineage negative (Lin-). This population may then be enriched further for markers associated with self-renewal capacity and multi-potency.

To visualize haematopoietic progenitor cells *in-vivo* using confocal or multi-photon microscopy, cells must be fluorescently labeled. There exist a number of strategies through which this is possible. The experiments from which the data used in this letter was extracted transplant and fluorescently labeled multi-potent progenitor (MPP) cells into an irradiated recipient. Fluorescent haematopoietic cells are extracted from mice genetically modified to express a florescent protein in their haematopoietic cells. Such cells are then purified into a rare population of multi-potent progenitor cells using fluorescence activated cell sorting to identify their associated cell surface markers. Figure 1 shows a schematic of this process.

#### Mice

Recipient mice are lethally irradiated using two doses of 5.5Gy of $\gamma$-radiation three hours apart.

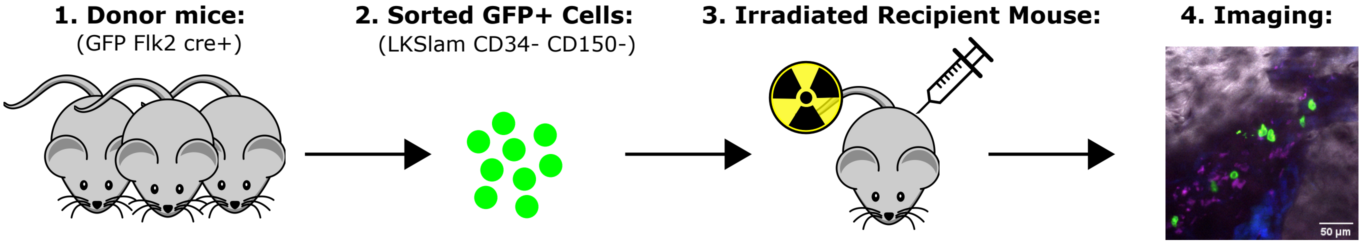


FIG. 1. *Experimental Setup* - Schematic showing the experimental setup used to generate the data used in this article. Bone marrow harvested from transgenic donor mice expressing a fluorophore protein in their haematopoietic cells is purified using fluorescence activated cell sorting (FACS) to produce a sub-set of bone marrow cells highly enriched for multi-potent progenitors (MPPs). These cells are then transplanted into syngeneic recipient mice and imaged using confocal microscopy two days after the transplantation.

#### Cell Sorting

The following set of cell surface markers were used to identify MPP cells: Lineage-, c-Kit+, Sca-1+, CD150-, CD48+. A '+' sign preceeding the marker name indicates that the cell population is enriched for that particular marker, likewise a '-' sign indicates depletion.

- Lin-: a cocktail of markers indicating terminally differentiated haematopoietic cells, namely CD3, CD4, CD8, B220, CD11b, Gr-1, and Ter119.
- Stem cell antigen Sca1+.
- Stem cell growth factor cKit+.
- CD48+ and CD150-.

#### Microscopy

Laser scanning confocal microscopy (LSCM) was used to record time-lapse images of the motion of MPP cells observed at various fixed positions within the calvarium bone marrow. Imaging commenced two days after transplantation was performed.

#### Data Acquisition

After the time-lapse images were recorded the central positions of the cell were extracted in a semi-automated fashion using IMARIS software.

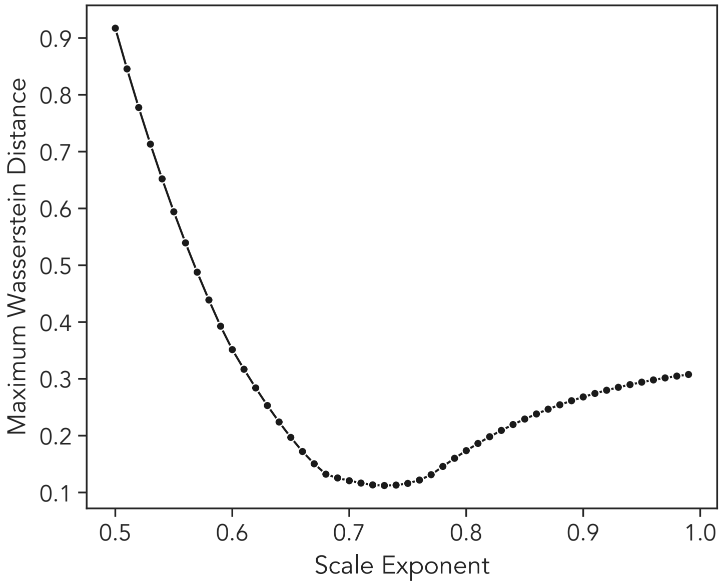


FIG. 2. *Optimization of scale exponent using Wasserstein distance* - The maximum value of the Wasserstein distance computed pairwise between the empirical cumulative distribution functions of the re-scaled variable $\eta = \delta x/\Delta^{\beta}$ where $\delta x = x(t+\Delta) - x(t)$, for each lag time $\Delta = \{3, 6, 15, 30, 60, 120\}$ mins used in the analysis. A clear minimum of this statistic is observed at a value of $\beta = 0.73$, which is the value used in figure 2(b) of the main text. All cells with track lengths greater than or equal to 120 mins were used in this analysis.

### DETAILS OF STATISTICAL ANALYSIS

#### Sensitivity Analysis of Curve Collapse

The value of the exponent $\beta$ was determined to be 0.73 using the following procedure. For a given value of the scale exponent $\beta'$, between each pair of empirical cumulative distribution functions for the re-scaled variable $\eta = \delta x/\Delta^{\beta'}$ computed for all lag times $\Delta = \{3, 6, 15, 30, 60, 120\}$ (mins) used in the analysis; we calculate the Wasserstein distance $l(u, v)$. The Wasserstein distance for two random variables with CDFs $U$ and $V$ is given by

$$l(u, v) = \int_{-\infty}^{\infty} |U - V| \, \mathrm{d}u\mathrm{d}v. \tag{1}$$

This was done for values of $\beta'$ between 0.5 and 1. The value of the exponent $\beta$ is chosen as the one which minimizes the maximum value of $l(u, v)$ between the distributions. The result is displayed in figure 2, from which a clear minimum at $\beta = 0.73$ can be observed, which is the value used to produce the curve collapse in figure 2(b) of the main text.

#### Quantification of Non-Gaussianity

Following on from the method of [1, 2] we attempt to quantify the non-Gaussian nature of the MPP displacement distributions over several lag times $\Delta$. To do this, for lag-times $\Delta = \{3, 6, 15, 30, 60, 120\}$ mins we plot distributions of the lagged x-displacement $\delta x = x(t + \Delta) - x(t)$ for all values $t \in [0, T - \Delta]$ where $T$ is the temporal length of a given track. We include all tracks long enough to produce at least one value of $\delta x$ for the largest lag time $\Delta_{max} = 120$ mins. We then, using maximum likelihood estimation(MLE) (implemented using SciPy [3]) estimated the parameters of a generalized Gaussian distribution

$$f(\delta x; \gamma, \sigma) = \frac{\gamma}{2\sigma^2\Gamma(1/\gamma)} \exp(|\delta x/\sigma|^{\gamma}) \tag{2}$$

where $\sigma$ is the scale parameter of the distribution, and $\Gamma(x)$ the Gamma function, required for proper normalisation of the distribution. The shape parameter $\gamma$ dictates the tailedness of the distribution - the greater it's value the

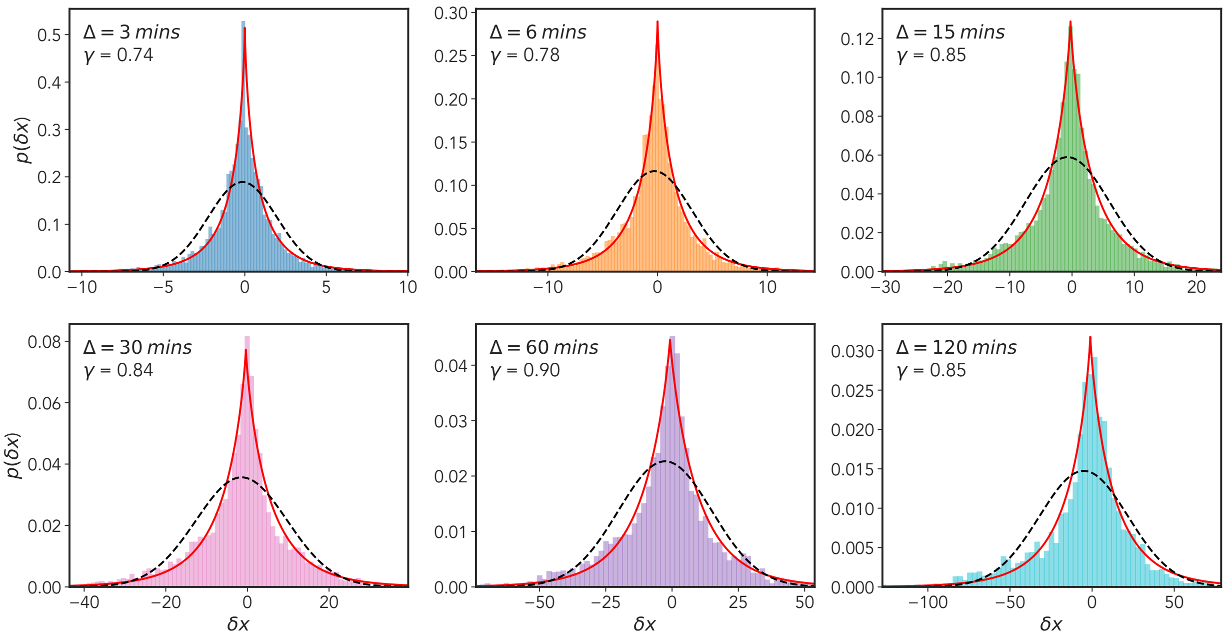


FIG. 3. *Non-Gaussian displacement distributions* - x-displacement distributions for six different lag times $\Delta = \{3, 6, 15, 30, 60, 120\}$ mins, where $\delta x = x(t+\Delta) - x(t)$ for all time points $t \in [0, T-\Delta]$ where $T$ is the length of the track. Cells included within this analysis were all those of track length greater than or equal to 120 mins. We show a maximum likelihood fit for both a generalized Gaussian distribution (red line) and a standard Gaussian (broken black line). The shape parameter $\gamma$ for the generalized Gaussian fit is shown along with the lag time $\Delta$ in the top left of each plot. In each case we see that the generalized Gaussian provides a superior fit.

more likely there is to be larger displacements. For $\gamma = 2$ the standard normal distribution is recovered, for $\gamma = 1$ a Laplacian is realized. The results of this analysis are shown in figure 3, with the solid red line representing the MLE fits. The fit for the shape parameter $\gamma$ reveals a spectrum of values all less than one, with a slight increasing trend over time. We also include the maximum likelihood estimates for a Gaussian distribution, plotted as a dashed black curve. Upon inspection, it is clear that the Gaussian distribution provides a poor fit at all values of lag time $\Delta$, while the generalized Gaussian, through incorporation of a shape parameter $\gamma$, arguably provides the minimal extension necessary required to provide a good fit.

Further to this, we also computed the non-Gaussianity parameter [4]

$$G(\Delta) = \frac{d}{d+2} \times \frac{\langle \overline{\delta^4(\Delta)} \rangle}{\langle \overline{\delta^2(\Delta)} \rangle^2} - 1 \tag{3}$$

where

$$\overline{\delta^4(\Delta)} = \frac{1}{T-\Delta} \int_0^{T-\Delta} (\mathbf{r}(t+\Delta) - \mathbf{r}(t))^4 \, \mathrm{d}t \tag{4}$$

and $\overline{\delta^2(\Delta)}$ is the TAMSD. This quantity is effectively a time-averaged version of the excess kurtosis of a probability distribution, with $G = 0$ for a Gaussian and $G > 0$ for a lepo-kurtotic fat tailed distribution - for example, a Laplacian. We display the results of this analysis, computed for the data used in figure 2(a) of the main text, in figure 4. We see that for small lag-times there is a pronounced departure from the expected result for a Gaussian distribution, $G(\Delta)$ then decays to approximately zero. Figure 4 when taken together with the results presented in figure 3, provides sufficient evidence to demonstrate the non-Gaussian nature of the underlying dynamical process generating the MPP motion. A notable point, is that the convergence of $G(\Delta)$ to the Gaussian value of $G = 0$ occurs prior to the cross over to Fickian diffusion identified within the main text. There has been substantial interest in the converse case, so-called "non-Gaussian yet Fickian diffusion". It is noted in [5] that slowly varying, heterogeneous fluctuations can lead to non-Gaussian displacement distributions, with a time-scale which persists to a time comparable to the cross-over to

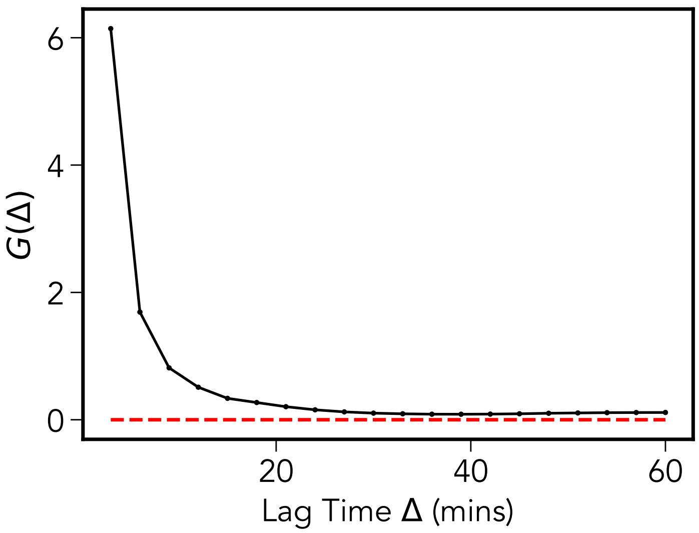


FIG. 4. *Non-Gaussianity Parameter* - the non-Gaussianity parameter $G(\Delta)$ shown for lag-times of upto one hour. At short times the departure from Gaussian behavior is marked. Longer times show a relaxation to the expected result for Gaussian processes $G = 0$.

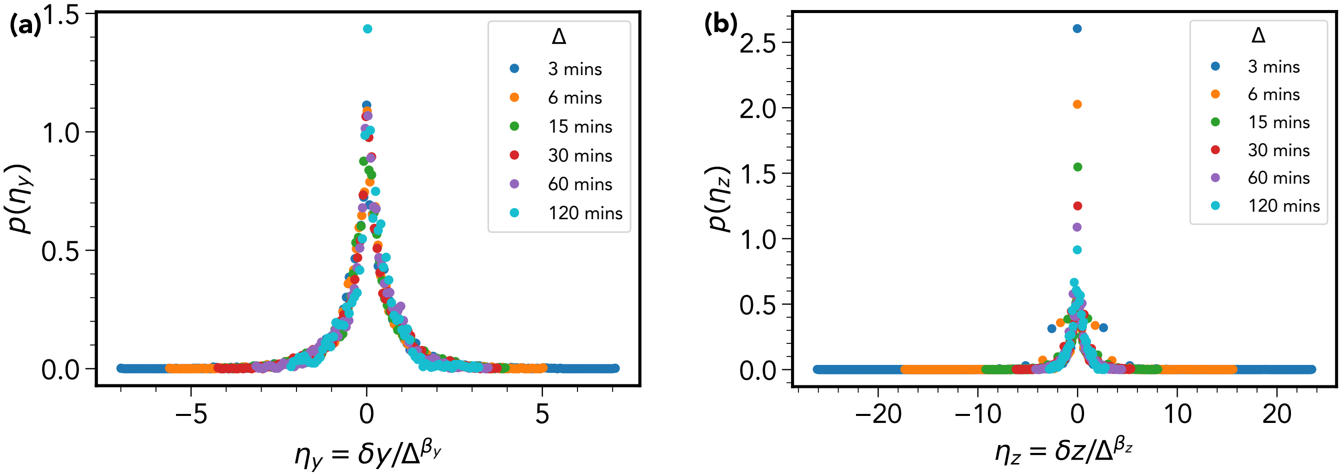


FIG. 5. *Curve collapse for y and z displacement distributions* - (a) A repeat of the figure 2(b) of the main text for the re-scaled y-displacement $\eta_y = \delta y/\Delta^{\beta_y}$, where we have determined $\beta = 0.76$. Figure (b) shows a the same plot for the z-displacement where this time $\beta_z = 0.59$. The $\eta_y$ distributions reproduce well the result in figure 2(b) of the main text, however the same cannot be said of $\eta_z$, which we attribute to sampling bias caused by a limited field of view in te z-direction.

Fickian diffusion. This is not the case for the MPP dynamics which appear to have a cross-over time to Gaussian behavior at $\tau_{gauss} \approx 30$ mins, where as the cross over to Fickian diffusion - as discussed in the main text is $\tau_{fick} \approx 150$ mins.

**Figure 2(b) for Y and Z Displacements and Confinement in the Z-Direction**

The curve collapse presented in figure 2(b) of the main text, used the re-scaled variable $\eta = \delta x/\Delta^{\beta}$. Where $\delta x$ represents the x-displacement, $\Delta$ the time-lag and $\beta$ the scale exponent determined via the procedure described above. In figure 5 we re-plot this for the y and z coordinates, this time using the notation $\eta_y = \delta y/\Delta^{\beta_y}$ and likewise for the

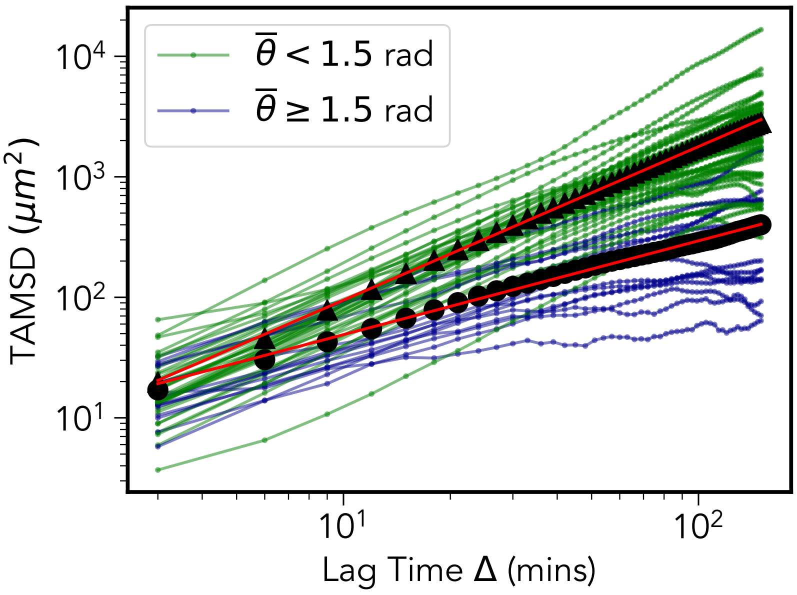


FIG. 6. *Time-Averaged Mean Square Displacement* - We label the TAMSD of more ($\theta < 1.5$) and less ($\theta \geq 1.5$) persistent cells with green and blue respectively. The corresponding averages over all cells within the more/less persistent groups are shown by the black triangles/circles, along with least-squares fit lines in red which yield exponents of 1.27 for the $\overline{\theta} < 1.5$ group, and 0.78 for $\overline{\theta} \geq 1.5$.

z-coordinate.

The distributions for the y-coordinate are extremely similar to that observed for $\delta x$ with $\beta_y$ this being determined to be 0.76. For the z displacement, the result was less convincing, the scale exponent $\beta_z$ was calculated to be 0.59. The reason for this anomaly is likely due to the fact that motion in the z-direction appears to be confined. This is an experimental artefact occurring due to the fact that there is a limited field of view in the z direction. Light from the confocal microscope is only able to penetrate down to fixed depth within the calvarium, therefore cells possessing a high degree of motility along z-axis will leave the field of view early on in the imaging window.

Therefore, longer tracks will belong to one of two categories; 1) motile cells which by chance happen to be moving pre-dominantly in the x-y plane, and 2) less motile cells unlikely to be able to leave the field of view during the observation time. We demonstrate this effect in figures 6 and 7. Dividing the MPP population into two categories based on the trajectory averaged turn angle $\overline{\theta}$ demonstrates that the TAMSD in figure 2(a) of the main text is dominated by a group of 37 (of 51) persistent cells ($\overline{\theta} < 1.5$) which - as demonstrated in figure 7(a) remain largely confined to motion within the x-y plane during the observation window. This may be contrasted with the group of 14 less persistent cells whose mean square displacement is impaired in all directions as shown in figure 7(b).

**Heterogeneity**

Heterogeneity has been implicated as an important factor contributing to the anomalous statistical behavior in populations of motile cells [4, 6, 7]. A number of theoretical and empirical studies have emerged demonstrating that the canonical indicators of super-diffusion; namely a scaling of the mean square displacement $\sim t^{\alpha}$ with $\alpha > 1$ and non-Gaussian displacement distributions can originate from simple models of heterogeneous persistent cell motion [6–9].

In particular, mathematical models incorporating heterogeneity have found particular use in explaining the so-called “Brownian yet non-Gaussian diffusion” in systems displaying a normal linear scaling of the mean square displacement with time, while having non-Gaussian step width distributions [5, 10–13]. It is noted [5] that this is facilitated by the lack of a strict separation of time-scales between the slowly varying heterogeneity and the onset of Fickian diffusion.

More pertinently, heterogeneity has also been postulated as a putative mechanism for super-diffusivity over long times in experimental tracking of ensembles of motile cells [6–9, 14–16]. This stands as an alternative to the Lévy walk model, which postulates a power law step length distribution, with uniformly distributed turn angles between

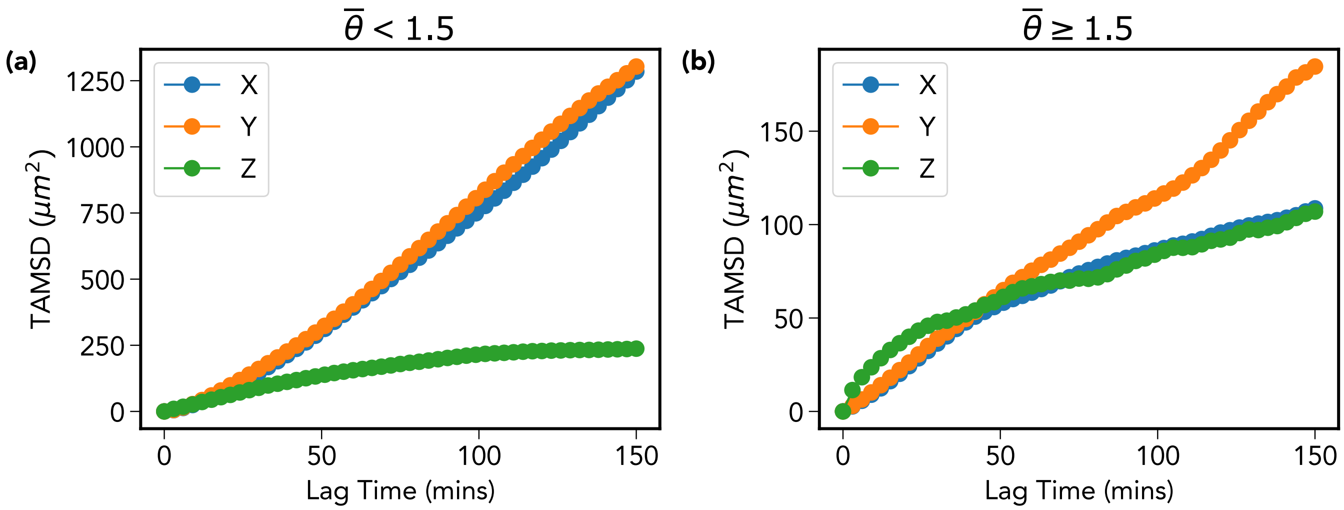


FIG. 7. *Confinement in the z-direction* - For all cells used in the TAMSD calculation there exists the appearance of a confinement effect in the z-direction due to the limited field of view in this direction. By dividing the cells into two populations based on their average turn angle, we can identify a group of (a) more persistent cells which by chance happen to moving in the x-y plane and (b) generally less motile cells which do not migrate sufficiently in any direction to leave the field of view.

steps [17–20]. This point is highlighted in a recent paper [9] where the authors compare a heterogeneous run-and-tumble model to a Lévy walk model as a mechanism for explaining the experimentally observed super-diffusive scaling of mouse fibroblast cells. The authors conclude that heterogeneous run and tumble motion provided a superior explanation for the observed super-diffusive scaling.

This work is one of a number of studies following recent trend towards more sophisticated statistical approaches aimed at making better use of the complex time series data obtained from single cell tracking experiments, often with the specific aim of accounting for the heterogeneity observed within experimental data. In particular, Bayesian methods [7, 21–23] and state space models such as the Hidden Markov Model [24–26] are increasingly utilized. Such methods may be used to complement traditional metrics such as the mean square displacement and step width distribution.

Heterogeneity in the dynamical behavior of motile cells has numerous causes. Some of these will be internal to the cell, reflecting variation in the transcriptional states of the underlying genome which pertain to cellular motility. Others will induced by environmental differences in the cells immediate surroundings, giving rise to a spatially dependent diffusion coefficient $D(x)$. This latter point is especially relevant in *in-vivo* settings such as the bone marrow cavity, which as a physical medium is known for it's complex heterogeneity. In *in-vitro* settings, environmental heterogeneity is directly controlled for, hence it is more natural to assume that each cell trajectory is generated by an identical probabilistic model. Following on from [8], it is therefore useful to outline two limiting cases which categorize heterogeneity within a cell population.

#### *Temporal Heterogeneity*

Temporal heterogeneity refers to statistical inhomogeneity with a single observed cell trajectory, is most typically manifested as stochastic alternation between different dynamical modes, for example, periods of inactivity where the cell centroid is effectively stationary, followed by bursts of motility. A recent paper on 2D super-diffusive behavior observed in a population on motile ameoba [1] noted this characteristic in their motion. During periods of relative inactivity the sampled positions of the cell centroid will not represent true translocations of the entire cell, but fluctuations of the centroid position caused by transient cell surface membrane fluctuations, which themselves are likely to be driven by re-arrangements of the underlying cellular cytoskeleton. Figure 8 (a) shows an example of an MPP trajectory along with it's corresponding displacement time-series in (b). Qualitatively we can see periods of very slow diffusion, followed by stretches of persistent motion. This characteristic will be problematic when constructing a model of cell locomotion which samples independently from the the empirical displacement and turn angle distributions, hence we it must be accounted for by any statistical model.

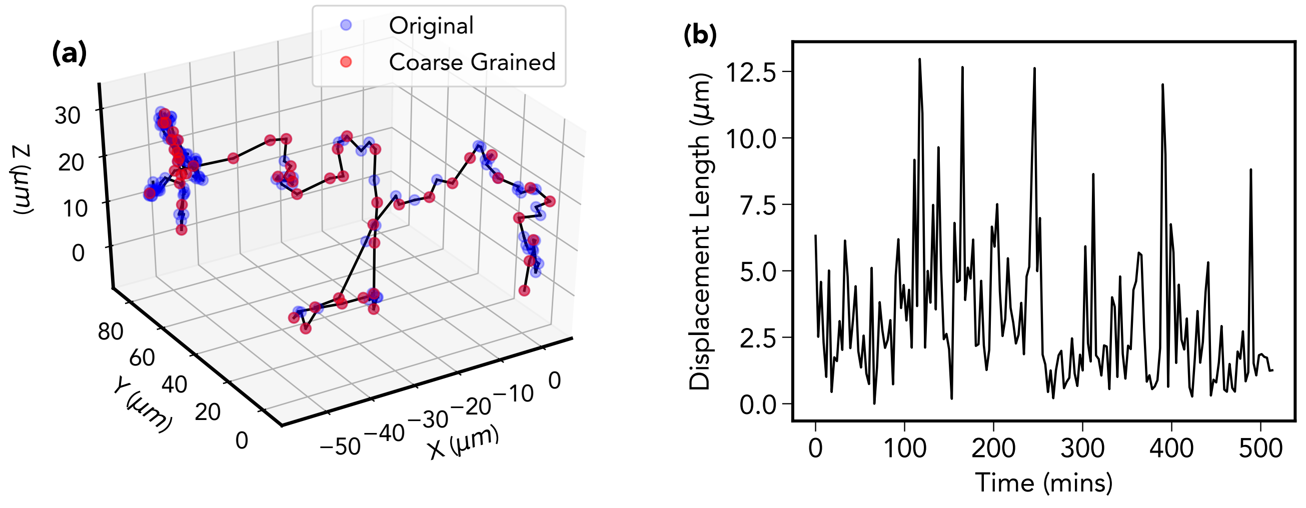


FIG. 8. *Temporal Heterogeneity* - (a) shows an example coarse-grained track (red) superimposed onto it's original (blue) in which each subsequent pair of points are separated by 3 minute intervals. Visually it appears as though periods of low motility and persistence are followed by more exploratory periods displaying persistent motion. During periods of lessened motility sampled displacements may not faithfully represented true translocation of entire cell; the coarse-graining procedure described within this SI lessens the effect of this heterogeneity. The displacement length time-series in shown for the same trajectory in (b).

*Cellular Heterogeneity*

Cellular heterogeneity reflects time independent, inter-cell variation in the statistical behavior of the cell trajectories. The origin of such variation is likely due to variation in the environment between each cells position within a given bone marrow cavity, on inter-mouse variation. This can be explained through variation in the mean run time $\bar{t}$, and variation in the mean turn angle $\bar{\theta}$ - this is shown in for the MPP trajectories in within the insets of figures 9(b)(c) and (d). Such inter-cell variation accounts for the significant spread in TAMSD curves shown in figure 2(a) of the main text, and therefore must be incorporated into our model to provide a faithful representation of the empirical data.

In the final section of the SI we demonstrate how the Bayesian inference method of [7] shows evidence of temporal and cellular heterogeneity within the ensemble of MPP trajectories.

### MODEL DESCRIPTION

#### Trajectory Coarse-Graining

An arbitrary CTRW may be defined through the specification of three distributions. The run length distribution $f(r)$, the run time distribution $g(\Delta t)$ and the turn angle distribution $g(\theta)$. Here we interpret a run to be undertaken at constant velocity, as opposed to a wait-time followed by a discontinuous jump. Many models following this prescription are also known as run-and-tumble (RTB) models, which typically have a completely Poissonian distributed run-time between subsequent uniformly distributed angular displacements. RTB models are heavily used in the description of the motion of Bacteria. The simplest case of such a model would have $r$ and $\tau$ related through a fixed speed $v$. For our data-set we will define a variable velocity model the using empirical cumulative distribution functions (ECDFs) computed directly from the data.

The empirical distributions are constructed by first applying a coarse-graining transformation to the trajectory. This involves including positions on the trajectory with at least $R$ microns of separation from their prior position, where $R$ is a threshold length-scale determined from the data. Given a track, which is a sequence of position vectors representing the cell centroid $\{\mathbf{r_t}\}$ observed at discrete times $t \in \{0, 1, 2, ..., T\}$ we extract a coarse-grained representation, which is an ordered subset of the original $\subseteq \{0, 1, 2, ..., T\}$ such that the initial time-point is always included and subsequent points are iteratively found as the first proceeding time point which satisfies the criterion $|\Delta\mathbf{r}_{t'} - \Delta\mathbf{r}_{t'-\Delta t}| > R$.

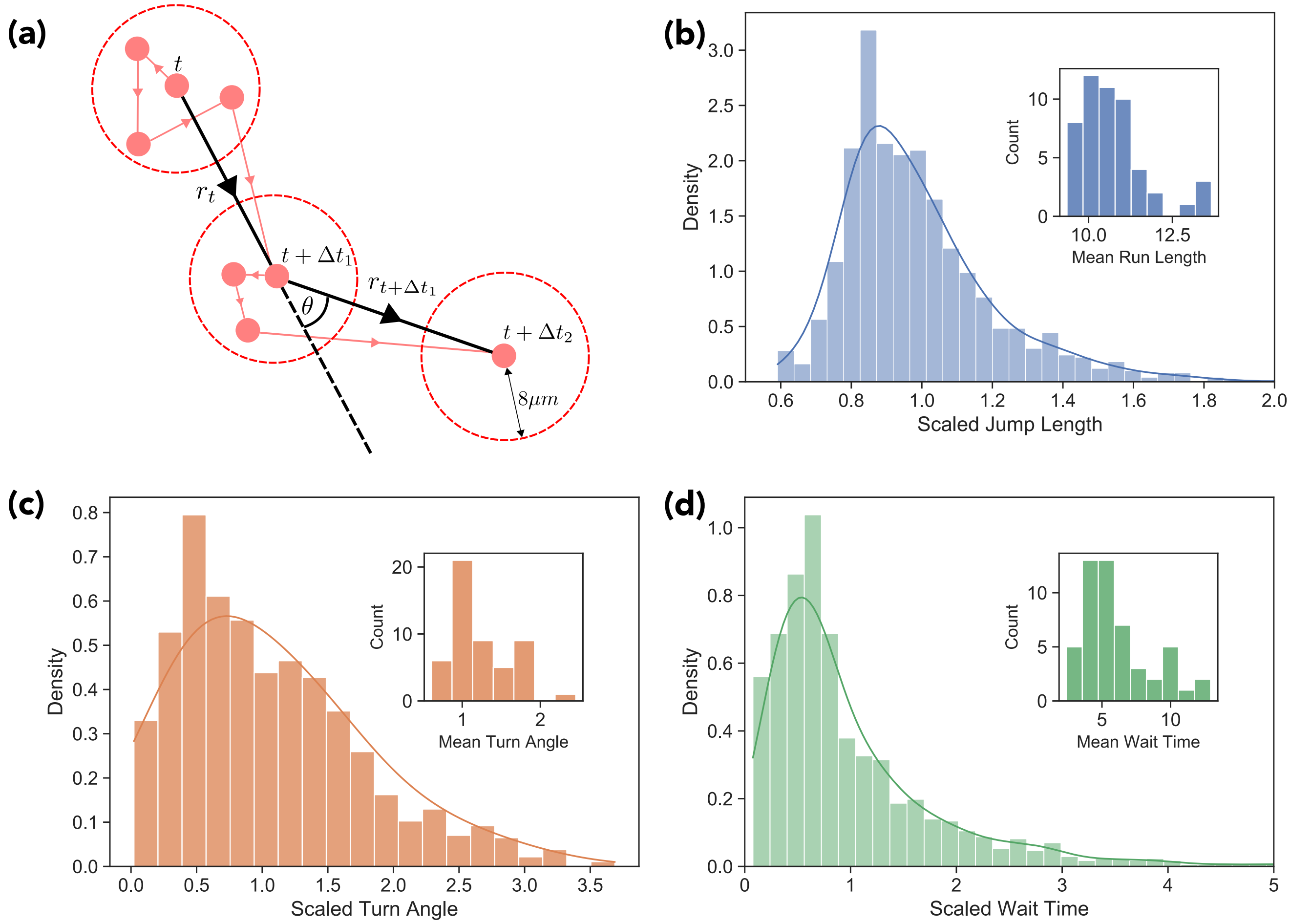


FIG. 9. *Coarse-grained trajectory* - schematic of three subsequent points along a coarse grained trajectory. The first point to have at least 6 microns displacement from the prior point is included in the coarse-grained trajectory. The displacement vector between two subsequent points is taken as the run vector with associated run-length $r$ and run-time $\Delta t$. Between two subsequent run vectors we calculate at turn angle $\theta_t$.

Between each pair of subsequent points on the track, we can therefore define a run length $r$ and an associated run time $\Delta t$, and a turn angle $\theta$ as the polar angle between two subsequent run vectors

$$\cos\theta_t = \frac{\mathbf{r}_t \cdot \mathbf{r}_{t+\Delta t}}{|\mathbf{r}_t| \cdot |\mathbf{r}_{t+\Delta t}|}. \tag{5}$$

We show a schematic of this process in figure 9(a).

This is a heuristic procedure done to remove short-range, transient fluctuations unrelated to genuine movements of the entire cell body. Without this transformation, simple repeated sampling from the ECDFs will not re-produce the mean square displacement of the experimental trajectories. This implies that any adequate model must account for the temporal correlation within the trajectories. The Bayesian approach of Metzner et al. [7] provides an elegant way in which to do this, but the coarse-graining procedure also suffices and has the advantage of that is has a very simple implementation; albeit while losing information pertaining to short-range movements of the cell body.

The threshold value $R$ was determined by minimizing the root sum of the squared error between the simulated and experimental EA-TAMSD.

#### Empirical Distributions

Using the coarse-grained tracks we are able to construct empirical distributions representing the run-length, run-time, and turn angle distributions required to specify the run and tumble model. However, as previously demonstrated, the MPP trajectories display significant cellular heterogeneity. This observation must therefore be incorporated into our model to faithfully reproduce the experimental statistical analysis. We do this by re-scaling each trajectory's run lengths, turn angles, and wait times by their corresponding mean values $\overline{r}, \overline{\theta}, \overline{\Delta t}$ and aggregating into dimensionless scaled run-length $f(r/\overline{\theta})$, scaled turn angle $g(\theta/\overline{\theta})$ and scaled run-time $h(\Delta t/\overline{\Delta t})$ empirical distribution functions (ECDFs). Histogram plots of the model distributions are shown in Fig. 9(b)(c)(d).

#### Model Algorithm

For each simulated cell trajectory we initially assign an empirical mean run-length, run-time and turn angle from one the associated 51 MPP trajectories used in the analysis. The algorithm proceeds by drawing dimensionless samples from the re-scaled empirical distributions, for example $f(r/\overline{r})$, and then multiplying the result with the corresponding mean value $\overline{r}_i$ to obtain the actual value. We begin by drawing initial scaled run length and run-time from distributions $f(r/\overline{r}), h(\Delta t/\overline{\Delta t})$ respectively.

As our data analysis and model are three-dimensional, two angles are required to specify a given direction. Assuming a spherical co-ordinate system, we have defined the turn angle $\theta$ between two subsequent runs as the polar angle between the two associated run displacement vectors. We assume no chirality, therefore when updating the position of a simulated trajectory we assign a random azimuthal angle $\phi$.

For each simulated cell trajectory we iteratively generate samples of the three relevant random variables $(r, \theta, \Delta t)$ using inverse transform sampling and update the position of cell using the scheme described below, until the prescribed temporal track length has been generated. To produce figure 3 of the main text we used linear interpolation to produce a track of simulated positions separated by evenly spaced three minute intervals. This interpolated trajectory could then be used to calculate the TAMSD and compare the model to the empirical data.

#### Trajectory Position Update Scheme

Given a current position $\mathbf{x}_t$ and previous cell position $\mathbf{x}_{t-\Delta t'}$ we wish to obtain the updated position $\mathbf{x}_{t+\Delta t}$ after a run defined by run length $r$, turn angle $\theta$, and run time $\Delta t$. We define run displacement vector $\mathbf{r}_t = \mathbf{x}_t - \mathbf{x}_{t-\Delta t'}$ and associated unit vector $\hat{\mathbf{r}}_t = \mathbf{r}_t/|\mathbf{r}_t|$ which specifies the polar axis of a reference frame centred on the point $\mathbf{x}_t$. We may then define a plane perpendicular to this direction which contains the point

$$\mathbf{x}_\perp = \mathbf{x}_t + r\cos(\theta)\hat{\mathbf{r}}_t \tag{6}$$

using

$$\hat{\mathbf{r}}_t \cdot \mathbf{x} = 0. \tag{7}$$

By expanding in terms of Cartesian co-ordinates and re-arranging we may eliminate one co-ordinate and obtain two basis vectors for the plane. Using the Gram-Schmidt procedure we can then obtain two orthogonal unit basis vectors $(\mathbf{e}_1, \mathbf{e}_2)$. We can then compute the new position of the trajectory after the run via

$$\mathbf{x}_{t+1} = \mathbf{x}_\perp + r\sin(\theta)(\cos(\phi)\mathbf{e}_1 + \sin(\phi)\mathbf{e}_2) \tag{8}$$

for an azimuthal angle $\phi \in [0, 2\pi)$ drawn from a uniform distribution. This procedure is iterated until the required temporal track length is generated, which in figure 3 of the main text is 30000 minutes per track.

#### Model Displacement Distributions

Figure 3. of the main text demonstrates that our model reproduces the scaling of the TAMSD. However, it is also informative to ask to what extent the model reproduces the non-Gaussian displacement distributions observed in the experimental data. In figure 10 we show a reproduction of figure 3, the distributions of the variable $\delta x = x(t+\Delta) - x(t)$

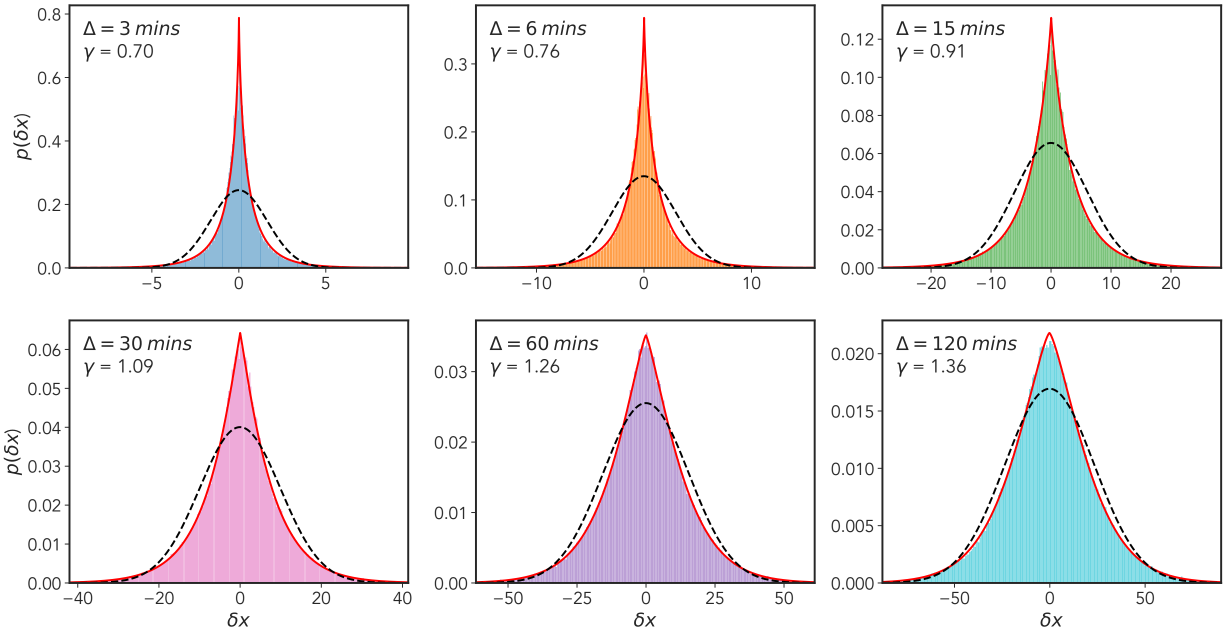


FIG. 10. *Model x-displacement distributions* - reproduction of figure 3 for the simulated data. As in figure 3 we show $x$-displacement distributions for six different lag times $\Delta = \{3, 6, 15, 30, 60, 120\}$ mins, where $\delta x = x(t+\Delta) - x(t)$ for all time points $t \in [0, T-\Delta]$ where $T$ is the length of the track. Each plot shows a maximum likelihood fit for both a generalized Gaussian distribution (red line) and a standard Gaussian (broken black line). The shape parameter $\gamma$ for the generalized Gaussian fit is shown along with the lag time $\Delta$ in the top left of each plot. Again in each case we see that the generalized Gaussian provides a superior fit, however, unlike the experimental data the Gaussian provides a progressively better fit with increasing lag time $\Delta$.

for increasing lag times $\Delta$. Interestingly, although the non-Gaussian form is still present - as reflected in the maximum likelihood fits - the distributions become progressively more Gaussian with increasing lag time. This trend does not occur to such an extent in the experimental data, the distribution $p(\delta x; \Delta_{max} = 120 \text{ mins})$ for the experimental data is still distinctly non-Gaussian with a shape parameter $\gamma < 1$. One possible explanation for this discrepancy is that our coarse graining procedure has significantly reduced the likelihood of observing short length displacements at longer lag times. In other words, we have smoothed out the effects of temporal heterogeneity within the track - reducing the contribution of the short-length cell centroid fluctuations observed for a given cell during a period of stationarity.

### ANALYSIS OF HETEROGENEITY USING SUPERSTATISTICAL BAYESIAN METHOD

#### Notation

In this section we use the following notation; we denote a collection of random variables $\{X_0, X_1, ..., X_T\}$ representing a discrete time stochastic process as $X_{0:T}$ and the corresponding observed values of this process in lower case notation as $x_{0:T}$. For continuous random variables we write the probability density of observing a given realization of the process as

$$p(x_{1:T})\mathrm{d}x_{1:T} = \Pr\big(X_1 \in (x_1, x_1 + \mathrm{d}x_1)\ ,\ \ldots\ldots,\ X_T \in (x_T, x_T + \mathrm{d}x_T)\big) \tag{9}$$

#### Model Specification

To demonstrate and quantify the nature of the heterogeneity present within the MPP trajectories we implement the Bayesian procedure of [7] in which cell motion is described hierarchically using a super-statistical model. In this context, the term super-statistics refers to models in which a low-level process representing the dynamics over short spatio-temporal scales, has slowly varying intensive parameters (for example temperature ) controlled by a high-level process [27, 28]. It has been demonstrated that the super-position of these two (or more) statistics can lead to systems with stationary states characterized by non-Gaussian fat-tailed probability distributions [29]. The method of [7], while inspired by this idea, is not completely faithful to it as there is not a strict separation of time-scales between the high level process. To account for rapid changes in motility parameters, regime changes where the cells transitions from a period of inactivity to higher motility, manifested by allowing for abrupt discontinuous jumps in parameter values are allowed. Specifically, Metzner et. al. describe cell motion using a discrete-time persistent random walk, or AR-1 process

$$\mathbf{v}_t = q_t \mathbf{v}_{t-1} + a_t \mathbf{n}_t \tag{10}$$

with time varying persistence $q_t$ and activity $a_t$ parameters. These parameters are viewed as random variables evolving according to a time-independent high-level process. This process is modeled through a transform $K$ of the parameter probability distribution.

### Bayesian Inference Algorithm

The inference algorithm is based on one of the most extensively used mathematical models for analysis of time series with time dependent states - the Hidden Markov Models (HMM) [30]. HMMs consider an observed discrete time-series as a collection of random variables $Y_{1:T} = \{Y_1, Y_2, ..., Y_T\}$ indexed by a time index $t$ whose values are conditionally independent when given the value of a hidden state $X_t$. The correlation structure of the observed time-series is encoded in the hidden state variables as a Markov process.

In our case, the observed variables correspond to the velocity data $\mathbf{v}_{1:T}$, and the hidden state the time-varying parameters $\boldsymbol{\theta}_t = (a_t, q_t)$. We wish to estimate the value of a latent parameter $\boldsymbol{\theta}_t = (a_t, q_t)$ given the entire observed time-series $\mathbf{v}_{1:T}$. Therefore, the inference has a naturally Bayesian interpretation, and is implemented using the forward-backward algorithm for HMMs [30, 31]. In the description of the forward-backward algorithm for simplicity we assume that the probability of a given observation depends only on the current value of the parameter, not on past data points. However, it is readily extended to auto-regressive cases such as that considered by Metzner et al. [7, 32, 33].

As with other Bayesian methods the parameters $\boldsymbol{\theta}_t = (a_t, q_t)$ are viewed as random variables, and the data-set of observed velocities $\mathbf{v}_{1:T}$ are interpreted as fixed. Bayes theorem is used to compute their posterior distribution through multiplication of the prior $p(\boldsymbol{\theta}_t)$ - representing previous knowledge of the parameter distribution - with a factorizable likelihood function $p(\mathbf{v}_{1:T}|\boldsymbol{\theta}_t) = \prod_{t=0}^{T} L_t$; where $L_t := p(\mathbf{v}_t|\boldsymbol{\theta}_t)$ is the one-step likelihood function. The normalization factor $p(\mathbf{v}_{1:t})$ is referred to as the model evidence and represents the relative likelihood that the data-set was generated by the model

$$p(\boldsymbol{\theta}_t|\mathbf{v}_{1:T}) = \frac{p(\mathbf{v}_{1:T}|\boldsymbol{\theta}_t)\; p(\boldsymbol{\theta}_t)}{p(\mathbf{v}_{1:T})} \tag{11}$$

for which it is possible to re-write as

$$p(\boldsymbol{\theta}_t|\mathbf{v}_{1:T}) = \frac{p(\mathbf{v}_{1:t}, \mathbf{v}_{t+1:T}|\boldsymbol{\theta}_t)\; p(\boldsymbol{\theta}_t)}{p(\mathbf{v}_{1:T})} \tag{12}$$

$$= \frac{p(\mathbf{v}_{1:t}, \boldsymbol{\theta}_t)\; p(\mathbf{v}_{t+1:T}|\boldsymbol{\theta}_t)}{p(\mathbf{v}_{1:T})} \tag{13}$$

Despite the somewhat cumbersome notation, the forward-backward algorithm results from repeated application of the chain rule for joint probability distributions and the conditional independence relations assumed by the model, in our case Markovian parameter dynamics. Denoting the state space of possible parameter values as $\Theta$

$$p(\boldsymbol{\theta}_t, \mathbf{v}_{1:t}) = \int_{\boldsymbol{\theta}\in\Theta} p(\boldsymbol{\theta}_t, \boldsymbol{\theta}_{t-1}, \mathbf{v}_{1:t})\, \mathrm{d}\boldsymbol{\theta}_{t-1} \tag{14}$$

$$= \int_{\boldsymbol{\theta}\in\Theta} p(\mathbf{v}_t, \mathbf{v}_{1:t-1}, \boldsymbol{\theta}_t, \boldsymbol{\theta}_{t-1})\, \mathrm{d}\boldsymbol{\theta}_{t-1} \tag{15}$$

$$= \int_{\boldsymbol{\theta}\in\Theta} p(\mathbf{v}_t|\boldsymbol{\theta}_t) p(\mathbf{v}_{1:t-1}, \boldsymbol{\theta}_t, \boldsymbol{\theta}_{t-1})\, \mathrm{d}\boldsymbol{\theta}_{t-1} \tag{16}$$

$$= \int_{\boldsymbol{\theta}\in\Theta} p(\mathbf{v}_t|\boldsymbol{\theta}_t) p(\mathbf{v}_{1:t-1}, \boldsymbol{\theta}_{t-1}) p(\boldsymbol{\theta}_t|\boldsymbol{\theta}_{t-1})\, \mathrm{d}\boldsymbol{\theta}_{t-1} \tag{17}$$

recognizing that the term $p(\mathbf{v}_{t-1}, \boldsymbol{\theta}_{t-1})$ is the joint parameter and data distribution for the previous time point $\alpha_{t-1}(\boldsymbol{\theta}_{t-1})$. The function $\alpha$ therefore obeys the following recursion relation

$$\alpha_t(\boldsymbol{\theta}_t) = p(\mathbf{v}_t|\boldsymbol{\theta}_t) \int p(\boldsymbol{\theta}_t|\boldsymbol{\theta}_{t-1})\alpha_{t-1}(\boldsymbol{\theta}_{t-1})\, \mathrm{d}\boldsymbol{\theta}_{t-1} \tag{18}$$

$$= L_t\, K^F \big[\alpha_{t-1}(\boldsymbol{\theta}_{t-1})\big] \tag{19}$$

A similar result may be obtained for the backward recursion for $\beta_t(\boldsymbol{\theta}_t)$ (see [31] for details)

$$\beta_t(\boldsymbol{\theta}_t) = \int_{\boldsymbol{\theta}\in\Theta} p(\boldsymbol{\theta}_{t+1}|\boldsymbol{\theta}_t)p(\mathbf{v}_{t+1}|\boldsymbol{\theta}_{t+1})\beta_{t+1}(\boldsymbol{\theta}_{t+1})\, \mathrm{d}\boldsymbol{\theta}_{t+1}$$
$$= K^B\big[L_{t+1}\beta_{t+1}\big]$$

returning to equation 12 we are now able to write in the form

$$\text{Posterior} \propto \text{Likelihood} \times \text{Prior}$$
$$= L_t \cdot K^F\big[\alpha_{t-1}\big]K^B\big[L_{t+1}\beta_{t+1}\big]$$
$$= L_t \cdot Pr_t^F \cdot Pr_t^B$$

where we have represented the integrals over the parameter space as integral transforms $K^F$ and $K^B$ with kernels $p(\boldsymbol{\theta}_t|\boldsymbol{\theta}_{t-1})$ and $p(\boldsymbol{\theta}_{t+1}|\boldsymbol{\theta}_t)$. If the parameter dynamics are time-reversible then $K^F = K^B$. We can therefore view the prior as the product of two independently obtained forward and backward priors $Pr^F$ and $Pr^B$ which have incorporated all information from the data in both directions of time converging onto to time $t$.

### Adapted Implementation of Metzner et. al.

Metzner at al. used a grid-based implementation of the above algorithm, in which the parameter state space $\Theta$ is discretized over a $200 \times 200$ rectangular grid. Starting from a flat prior the forward and backward recursions are run independently from time-points $t+2$ and $T$ respectively. The parameter bounds are $q_t \in (-1, 1)$ and $a_t \in (0, a_{max})$ where the limit $a_{max}$ is data-set dependent. Apart from the discrete approximation which facilitates computational efficiency and allows for a direct estimation of the model evidence; the main ingenuity of this technique is to generalize the integral transform $K$ of the forward and backward priors $\alpha$ and $\beta$ to allow for a more general class of transformations. In cases such as this, where there is no prior knowledge regarding the form of the parameter dynamics it is preferred to keep the transformation $K$ as general as possible, as to encapsulate the both gradual and abrupt possible parameter changes while not relying on a specific functional form. This process is shown graphically in figure 11. The likelihood function $L_t$ is approximated directly from the data and multiplied with priors. Following on from [7] we use a two step process to transform the parameters; firstly, to account for possible abrupt parameter changes, a minimum probability $p_{min}$ is assigned to each point of the grid. Secondly, to account for gradual changes in parameter values, we apply a uniform convolution to the parameter grid with box of radius $R$.

### Results of Bayesian Analysis

To fix the value of the hyper-parameters $p_{min}$ and $R$ controlling the high-level parameter transformation, we select the values of $p_{min}$ and $R$ which maximize the model evidence $p(\mathbf{v}_{1:T})$, evaluated by summing the (un-normalized) posterior density over all points of the grid. The model evidence provides a quantitative measure of the likelihood that the data was generated by the model. The discrete sum approximates the integral

$$p(\mathbf{v}_{1:T}|p_{min}, R) = \int_{\boldsymbol{\theta}\in\Theta} p(\boldsymbol{\theta}_{1:T}, \mathbf{v}_{1:T}|p_{min}, R) \approx \sum_{ij} p(\boldsymbol{\theta}_{ij}, \mathbf{v}_{1:T}|p_{min}, R) \cdot \Delta_{\boldsymbol{\theta}} \tag{20}$$

where we use the indices $ij$ to represent the points of the discretized parameter grid and $\Delta_{\boldsymbol{\theta}}$ represents the voxel size of the grid. We also note that the results presented herein are not overly sensitive to variation of these hyper-parameters, however, we use the model evidence maximum maximization criteria as means to fix their value. This is a form of what is known as empirical Bayes in the mathematical statistics literature.

In figure 12 we show the result of this analysis applied to a particular trajectory from the ensemble of MPP tracks. For each posterior $Po_t$ we compute the means of both the activity and persistence parameters, along with their 50

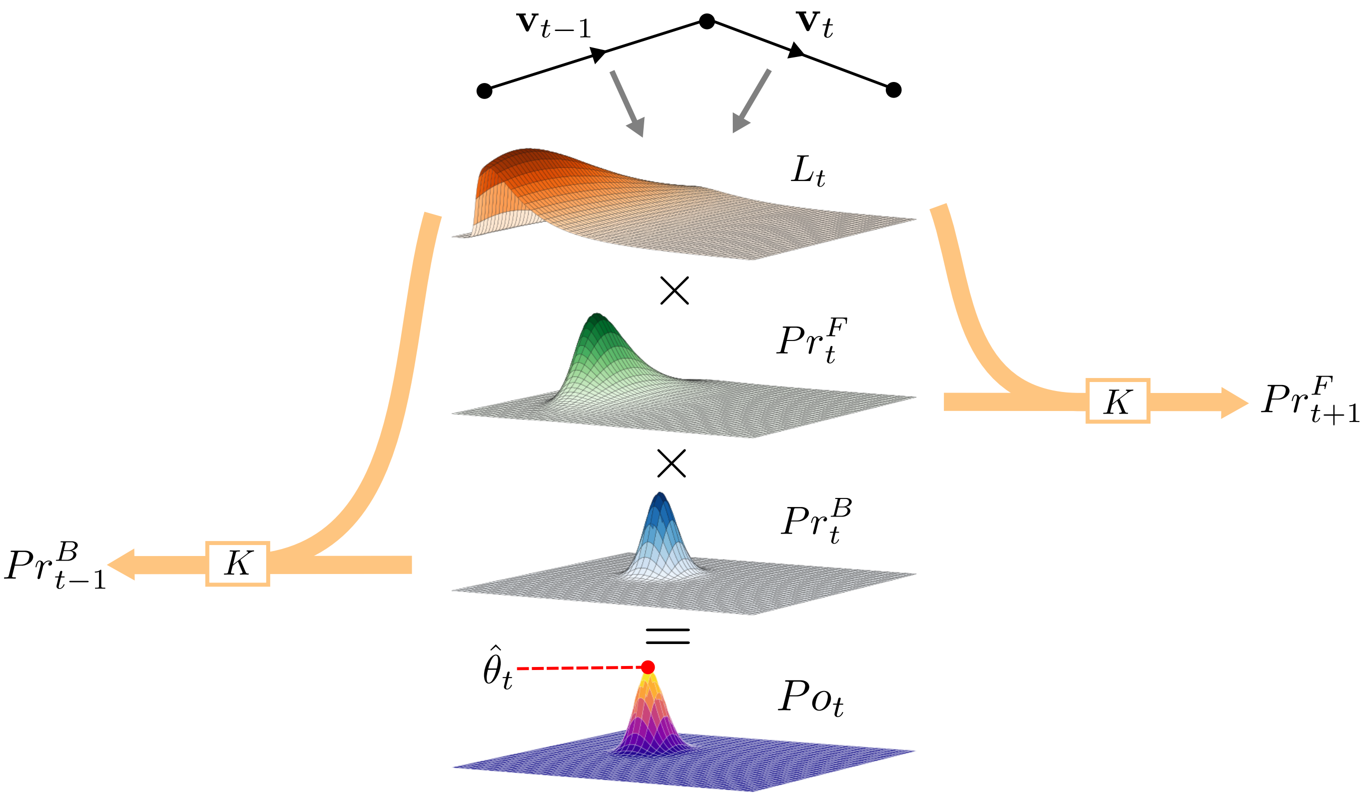


FIG. 11. *Schematic of superstatistical Bayesian method* - Given an experimental time-series and an observation likelihood function $L$, which here we assume to be of the form of the Gaussian AR-1 process described in the main text, it is possible to compute the posterior distribution of the time-varying activity $a_t$ and persistence $q_t$ parameters using a modified, discretized version of the forward-backward algorithm for hidden Markov models. The posterior parameter distribution is obtained through multiplication of the grids representing the likelihood $L_t$ and the forward and backward priors $Pr_t^F$ and $Pr_t^B$ obtained using the recursion relations presented in the main text. Point parameter estimates $\hat{\theta}_t$ can be obtained from this posterior, typically through the mean or the mode. The subsequent priors in the forward and backward direction are computed independently using the ad-hoc transformation $K$.

percent credible regions, shown in figures 12(c) and (d). In figure 12(a) we annotate a trajectory according to the inferred persistence parameter values. Visually, it is clearly seen that the algorithm is able to pick out periods of persistent motion interspersed with periods of low activity and persistence. The time-averaged posterior distribution shown in figure 12 (b) provides further insight into the motion of cell over the entire trajectory; for this particular track we see that there exists three distinct modes, one low activity anti-persistent, one low-activity persistent, and one high activity.

Repeating this analysis for the entire ensemble of MPP tracks, we observe a large spread in the values of the $a_t$ and $q_t$, shown in the histogram plot in figure 13.

---

*
†
[1] A. G. Cherstvy, O. Nagel, C. Beta, and R. Metzler, Non-Gaussianity, population heterogeneity, and transient superdiffusion in the spreading dynamics of amoeboid cells, Physical Chemistry Chemical Physics **20**, 23034 (2018).
[2] T. J. Lampo, S. Stylianidou, M. P. Backlund, P. A. Wiggins, and A. J. Spakowitz, Cytoplasmic RNA-Protein Particles Exhibit Non-Gaussian Subdiffusive Behavior, Biophysical Journal **112**, 532 (2017).
[3] P. Virtanen, R. Gommers, T. E. Oliphant, M. Haberland, T. Reddy, D. Cournapeau, E. Burovski, P. Peterson, W. Weckesser, J. Bright, S. J. van der Walt, M. Brett, J. Wilson, K. J. Millman, N. Mayorov, A. R. Nelson, E. Jones,

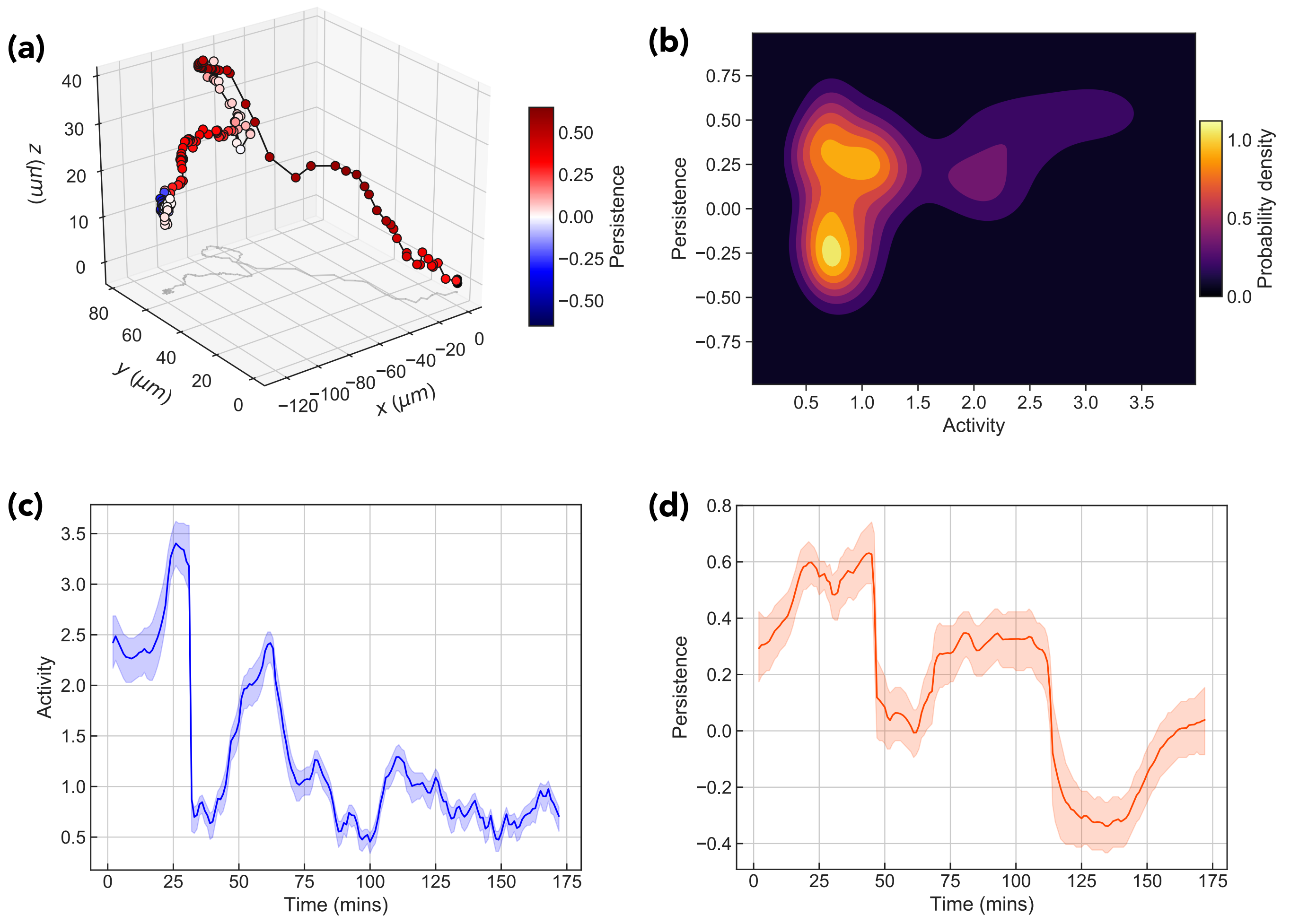


FIG. 12. *Results of Bayesian analysis for example trajectory* - (a) Shows an example MPP trajectory color coded according to the value of the persistence parameter $q$ at that time. Figure (b) shows the time-averaged mean posterior distribution. Figures (c) and (d) show the time evolution of the posterior mean of the activity parameter $a$ and persistence parameter $q$ respectively.

R. Kern, E. Larson, C. J. Carey, l. Polat, Y. Feng, E. W. Moore, J. VanderPlas, D. Laxalde, J. Perktold, R. Cimrman, I. Henriksen, E. A. Quintero, C. R. Harris, A. M. Archibald, A. H. Ribeiro, F. Pedregosa, P. van Mulbregt, A. Vijaykumar, A. P. Bardelli, A. Rothberg, A. Hilboll, A. Kloeckner, A. Scopatz, A. Lee, A. Rokem, C. N. Woods, C. Fulton, C. Masson, C. Häggström, C. Fitzgerald, D. A. Nicholson, D. R. Hagen, D. V. Pasechnik, E. Olivetti, E. Martin, E. Wieser, F. Silva, F. Lenders, F. Wilhelm, G. Young, G. A. Price, G. L. Ingold, G. E. Allen, G. R. Lee, H. Audren, I. Probst, J. P. Dietrich, J. Silterra, J. T. Webber, J. Slavič, J. Nothman, J. Buchner, J. Kulick, J. L. Schönberger, J. V. de Miranda Cardoso, J. Reimer, J. Harrington, J. L. C. Rodríguez, J. Nunez-Iglesias, J. Kuczynski, K. Tritz, M. Thoma, M. Newville, M. Kümmerer, M. Bolingbroke, M. Tartre, M. Pak, N. J. Smith, N. Nowaczyk, N. Shebanov, O. Pavlyk, P. A. Brodtkorb, P. Lee, R. T. McGibbon, R. Feldbauer, S. Lewis, S. Tygier, S. Sievert, S. Vigna, S. Peterson, S. More, T. Pudlik, T. Oshima, T. J. Pingel, T. P. Robitaille, T. Spura, T. R. Jones, T. Cera, T. Leslie, T. Zito, T. Krauss, U. Upadhyay, Y. O. Halchenko, and Y. Vázquez-Baeza, SciPy 1.0: fundamental algorithms for scientific computing in Python, Nature Methods **17**, 261 (2020), arXiv:1907.10121.

[4] R. Metzler, J.-H. Jeon, A. G. Cherstvy, and E. Barkai, Anomalous diffusion models and their properties: non-stationarity, non-ergodicity, and ageing at the centenary of single particle tracking, Phys. Chem. Chem. Phys. **16**, 24128 (2014).

[5] B. Wang, J. Kuo, S. C. Bae, and S. Granick, When Brownian diffusion is not Gaussian, Nature Materials **11**, 481 (2012).

[6] S. Hapca, J. W. Crawford, and I. M. Young, Anomalous diffusion of heterogeneous populations characterized by normal diffusion at the individual level, Journal of the Royal Society Interface **6**, 111 (2009).

[7] C. Metzner, C. Mark, J. Steinwachs, L. Lautscham, F. Stadler, and B. Fabry, Superstatistical analysis and modelling of heterogeneous random walks, Nature Communications 10.1038/ncomms8516 (2015).

[8] T. Kwon, O. S. Kwon, H. J. Cha, and B. J. Sung, Stochastic and Heterogeneous Cancer Cell Migration: Experiment and Theory, Scientific Reports **9**, 1 (2019).

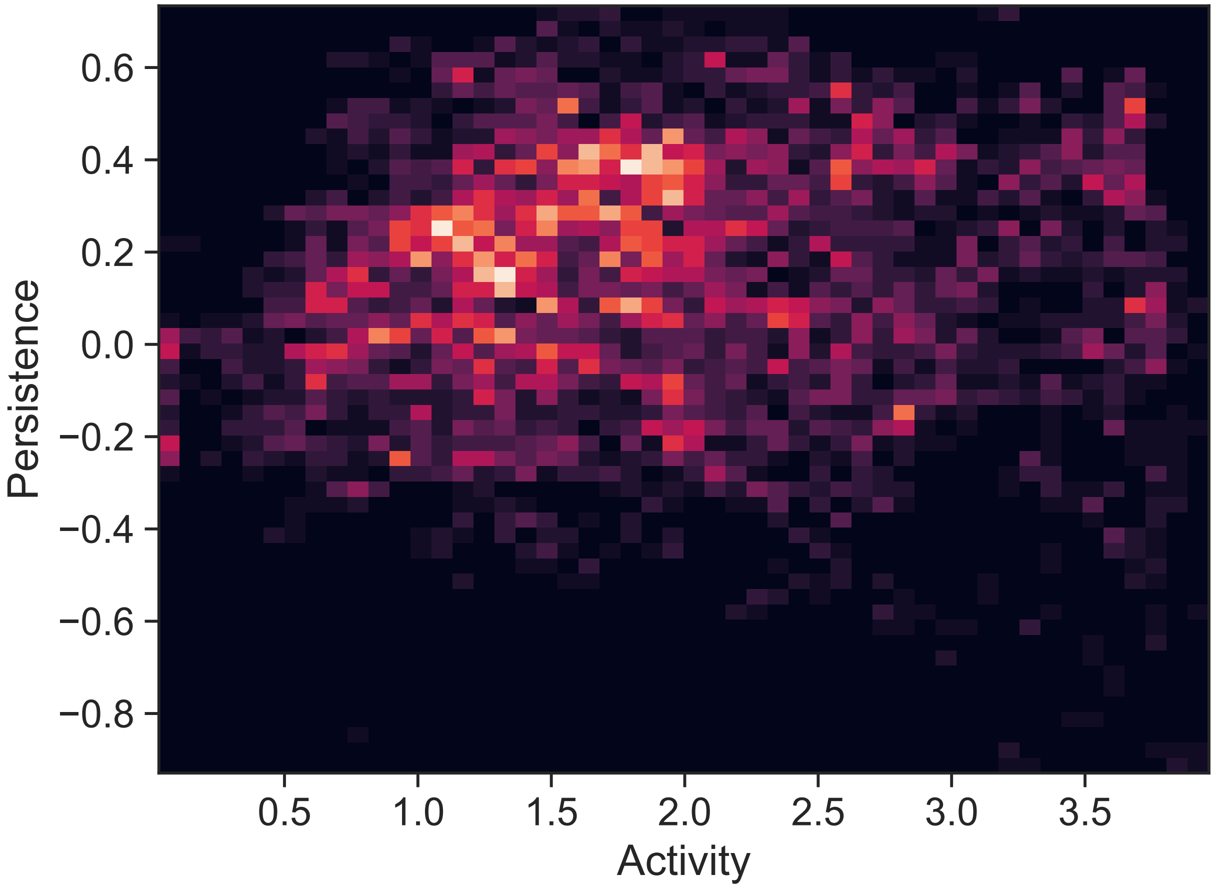


FIG. 13. *Results of Bayesian analysis for entire ensemble of MPPs* - 2D histogram plot of time varying activity and persistence parameters $a_t$ and $q_t$ collated over all MPP tracks.

[19] G. Ariel, A. Be'Er, and A. Reynolds, Chaotic Model for Lévy Walks in Swarming Bacteria, Physical Review Letters **118**, 1 (2017).
[20] S. Huda, B. Weigelin, K. Wolf, K. V. Tretiakov, K. Polev, G. Wilk, M. Iwasa, F. S. Emami, J. W. Narojczyk, M. Banaszak, S. Soh, D. Pilans, A. Vahid, M. Makurath, P. Friedl, G. G. Borisy, K. Kandere-Grzybowska, and B. A. Grzybowski, Lévy-like movement patterns of metastatic cancer cells revealed in microfabricated systems and implicated in vivo, Nature Communications **9**, 1 (2018).
[21] N. Monnier, S. M. Guo, M. Mori, J. He, P. Lénárt, and M. Bathe, Bayesian approach to MSD-based analysis of particle motion in live cells, Biophysical Journal **103**, 616 (2012).
[22] F. Persson, M. Lindén, C. Unoson, and J. Elf, Extracting intracellular diffusive states and transition rates from single-molecule tracking data, Nature Methods **10**, 265 (2013).
[23] C. Mark, C. Metzner, L. Lautscham, P. L. Strissel, R. Strick, and B. Fabry, Bayesian model selection for complex dynamic systems, Nature Communications **9**, 10.1038/s41467-018-04241-5 (2018).
[24] G. Rosser, A. G. Fletcher, D. A. Wilkinson, J. A. de Beyer, C. A. Yates, J. P. Armitage, P. K. Maini, and R. E. Baker, Novel Methods for Analysing Bacterial Tracks Reveal Persistence in Rhodobacter sphaeroides, PLoS Computational Biology **9**, 10.1371/journal.pcbi.1003276 (2013).
[25] Y. Matsuda, I. Hanasaki, R. Iwao, H. Yamaguchi, and T. Niimi, Estimation of diffusive states from single-particle trajectory in heterogeneous medium using machine-learning methods, Physical Chemistry Chemical Physics **20**, 24099 (2018).
[26] J. Janczura, M. Balcerek, K. Burnecki, A. Sabri, M. Weiss, and D. Krapf, Identifying heterogeneous diffusion states in the cytoplasm by a hidden Markov model, New Journal of Physics **23**, 10.1088/1367-2630/abf204 (2021).
[27] C. Beck and E. G. Cohen, Superstatistics, Physica A: Statistical Mechanics and its Applications **322**, 267 (2003), arXiv:0205097 [cond-mat].
[28] C. Beck, Generalized statistical mechanics for superstatistical systems, Philosophical Transactions of the Royal Society A: Mathematical, Physical and Engineering Sciences **369**, 453 (2011).
[29] C. Beck, Stretched exponentials from superstatistics, Physica A: Statistical Mechanics and its Applications **365**, 96 (2006), arXiv:0510841 [cond-mat].
[30] L. R. Rabiner, A Tutorial on Hidden Markov Models and Selected Applications in Speech Recognition, Proceedings of the IEEE **77**, 257 (1989).
[31] C. M. Bishop, *Pattern Recognition and Machine Learning* (Springer-Verlag New York, 2007).
[32] I. Stanculescu, C. K. Williams, and Y. Freer, Autoregressive Hidden Markov Models for the Early Detection of Neonatal Sepsis, IEEE Journal of Biomedical and Health Informatics **18**, 1560 (2014).
[33] C. Mark, C. Metzner, and B. Fabry, Bayesian inference of time varying parameters in autoregressive processes, arXiv , 1 (2014), arXiv:1405.1668.
